# Gut microbiome of an unindustrialized population have characteristic enrichment of SNPs in species and functions with the succession of seasons

**DOI:** 10.1101/418889

**Authors:** Jiyue Qin, Chongyang Tan, Kang Ning

## Abstract

Most studies investigating human gut microbiome dynamics are conducted in modern populations. However, unindustrialized populations are arguably better subjects in answering human-gut microbiome coevolution questions due to their lower exposure to antibiotics and higher dependence on natural resources. Hadza hunter-gatherers in Tanzania have been found to exhibit high biodiversity and seasonal patterns in their gut microbiome composition at family level, where some taxa disappear in one season and reappear at later time. However, such seasonal changes have previously been profiled only according to species abundances, with genome-level variant dynamics unexplored. As a result it is still elusive how microbial communities change at the genome-level under environmental pressures caused by seasonal changes. Here, a strain-level SNP analysis of Hadza gut metagenome is performed for 40 Hadza fecal samples collected in three seasons. First, we benchmarked three SNP calling tools based on simulated sequencing reads, and selected VarScan2 that has highest accuracy and sensitivity after a filtering step. Second, we applied VarScan2 on Hadza gut microbiome, with results showing that: with more SNP presented in wet season in general, eight prevalent species have significant SNP enrichments in wet season of which only three species have relatively high abundances. This indicates that SNP characteristics are independent of species abundances, and provides us a unique lens towards microbial community dynamics. Finally, we identify 83 genes with the most characteristic SNP distributions between wet season and dry season. Many of these genes are from *Ruminococcus obeum*, and mainly from metabolic pathways like carbon metabolism, pyruvate metabolism and glycolysis, as shown by KEGG annotation. This implies that the seasonal changes might indirectly impact the mutational patterns for specific species and functions for gut microbiome of an unindustrialized population, indicating the role of these variants in their adaptation to the changing environment and diets.

**Importance:** By analyzing the changes of SNP enrichments in different seasons, we have found that SNP characteristics are independent of species abundances, and could provide us a unique lens towards microbial community dynamics at the genomic level. Many of the genes in microbiome also presented characteristic SNP distributions between wet season and dry season, indicating the role of variants in specific species in their adaptation to the changing environment for an unindustrialized population.

## Introduction

Thanks to the advancement of next-generation sequencing and bioinformatics techniques, we could now reveal all genetic information for the entire microbial community. Human gut microbiome is important due to its critical role in metabolism, host nutrition, immune function, and central nervous system[1, 2]. While most studies investigating human gut microbiome dynamics are conducted in modern populations, unindustrialized populations, with their lower exposure to antibiotics and higher dependence on natural resources, are arguably better subjects in answering human-gut microbiome coevolution questions[3]. Recent studies about the gut microbiome of the Hadza hunter-gatherers have shed light on the dynamics and adaptive versatility of the microbiome to lifestyle changes. The Hadza gut microbial communities exhibit a high degree of biodiversity[4] and a pattern of seasonal cycling in gut microbiome composition, where some taxa (at the family level) disappear in one season and reappear at later timepoint[5].

However, such seasonal changes of the Hadza gut microbial communities are previously profiled only according to species abundances, with genome-level variant dynamics unexplored. It was still unclear how microbial communities change at the genome-level under environmental pressures caused by seasonal changes.

The genome-level variations of the Hadza gut microbial communities could be examined by SNP profile assessment. As genomic variations like SNPs can lead to changes in antibiotic resistance[6] or pathogenicity[7], which can indicate the response to selection pressures[8], it is interesting to perform high-resolution investigation into the Hadza gut microbiome. The pipeline for analyzing genomic variants devised by Schloissnig[9] have been applied to studies about microbiomes[10] and their association with human diseases[11].

Here, we performed a strain-level SNP analysis of Hadza gut metagenome to decipher the microbiome dynamics from the perspective of SNP enrichment. We first evaluated the SNP calling methods on metagenome datasets, and selected the best method, namely VarScan2, for SNP calling on Hadza gut metagenome. Then, we performed in-depth analysis of the SNP enrichments in species and functions along the seasonal shifts, and attempted to interpret the dynamic patterns for such enrichments.

## Results and Discussions

### Assessment of SNP-calling tools

To select the most suitable variant-calling tool for the strain-level SNP analysis, we first evaluated three representative tools (GATK[12], BCFtools[13], VarScan2[14]) based on their performance in terms of sensitivity and selectivity (see **Materials and Methods**). A SNP list containing 8,148 sites from 4 species residing in human gut (*Bacteroides caccae, Bacteroides fragilis, Bacteroides ovatus, Bacteroides vulgatus*) was used to generate a mutated reference genome set. BBMap was then applied to generate 1,000,000 paired-end simulated reads with the sequencing error: SNPrate=0.002, insrate=0.00002, delrate=0.00002. We then used Burrows-Wheeler Aligner(BWA) to index the original reference genome set and MEM algorithm to align the simulated reads with the reference. The resulting sam files were then sent through a pretreatment procedure, which includes a transversion to bam files via SAMtools[15] view, a sorting process via SAMtools sort and a process to remove PCR duplicates via Picard. Three tools were used to call SNP from those processed bam files. BCFtools, GATK, VarScan2 identified 5,989, 5,670 and 4,298 raw SNP sites, respectively. Comparing the identified SNP with actual SNP in the mutated reference genome set, we found that BCFtools shows highest sensitivity but lowest selectivity, while VarScan2 shows highest selectivity but lowest sensitivity (**Table 1**). After selecting the SNP sites whose sequencing depth exceeds 9, we find that VarScan2 shows better sensitivity and best selectivity (**Table 2**). Thus, we have chosen VarScan2 for the human metagenome SNP analysis. Additionally, we can conclude that this filtering step largely decreases the number of false-positive SNP calls, which indicates the importance of performing filtering with a suitable threshold to obtain reliable SNP for further analysis.

**Table 1.** Comparison of raw SNP identification results by the three tools

|  | # Mismatch | # False positive | # All | Sensitivity | Selectivity |
| --- | --- | --- | --- | --- | --- |
| BCFtools | 1 | 143 | 8,148 | 0.717 | 0.982 |
| GATK | 1 | 91 | 8,148 | 0.685 | 0.989 |
| VarScan2 | 0 | 1 | 8,148 | 0.527 | 1.000 |

**Table 2.** Comparison of filtered SNP identified by the three tools

|  | # Mismatch | # False positive | # All | Sensitivity | Selectivity | # Mismatch |
| --- | --- | --- | --- | --- | --- | --- |
| BCFtools | 819 | 0 | 5 | 8,148 | 0.101 | 0.999 |
| GATK | 704 | 0 | 23 | 8,148 | 0.086 | 0.997 |
| VarScan2 | 819 | 0 | 0 | 8,148 | 0.101 | 1 |

Though we have attempted to mimic the actual variation in human gut microbiome, there were still some limitations in these simulated datasets. For example, the simulated reads were based on four gut species which were less than the actual number of species in gut microbial communities, and the relative quantities of these species were not considered. However, based on such simplified simulated metagenomic datasets, the advantages of VarScan2 over the other two methods are still obvious.

### Community compositions and their seasonal changes for Hadza gut microbiome

To remove adapters and low-quality bases, we first applied Trimmomatic[16] to Hadza’s gut whole-genome-sequencing dataset (sample size: 40). The parameters used for Trimmomatic were SE-phred33 ILLUMINACLIP:TruSeq3-SE.fa:2:30:10 LEADING:5 TRAILING:5, which removes adapters and low-quality bases. MetaPhlAn2[17] was utilized to determine the bacterial species and their abundance in each sample. A total of 116 species, including 16 unclassified species, were identified. Hadza gut microbial community’s species composition is shown in **Figure 1**. We can see that the main species include *Faecalibacterium prausnitzii*, *Prevotella copri*, *Methanobrevibacter smithii*, *Eubacterium biforme, Treponema succinifaciens* which corresponds with the previous report about the high abundance of *Prevotella* and *Treponema*[4]. Although *Treponema* is famous for one species, namely *T. pallidum*, which causes syphilis, here we demonstrate that *Treponema* in Hadza gut is mainly *T.succinifaciens*, which plays a role in cellulose and xylose hydrolysis[18].

**Figure 1.**
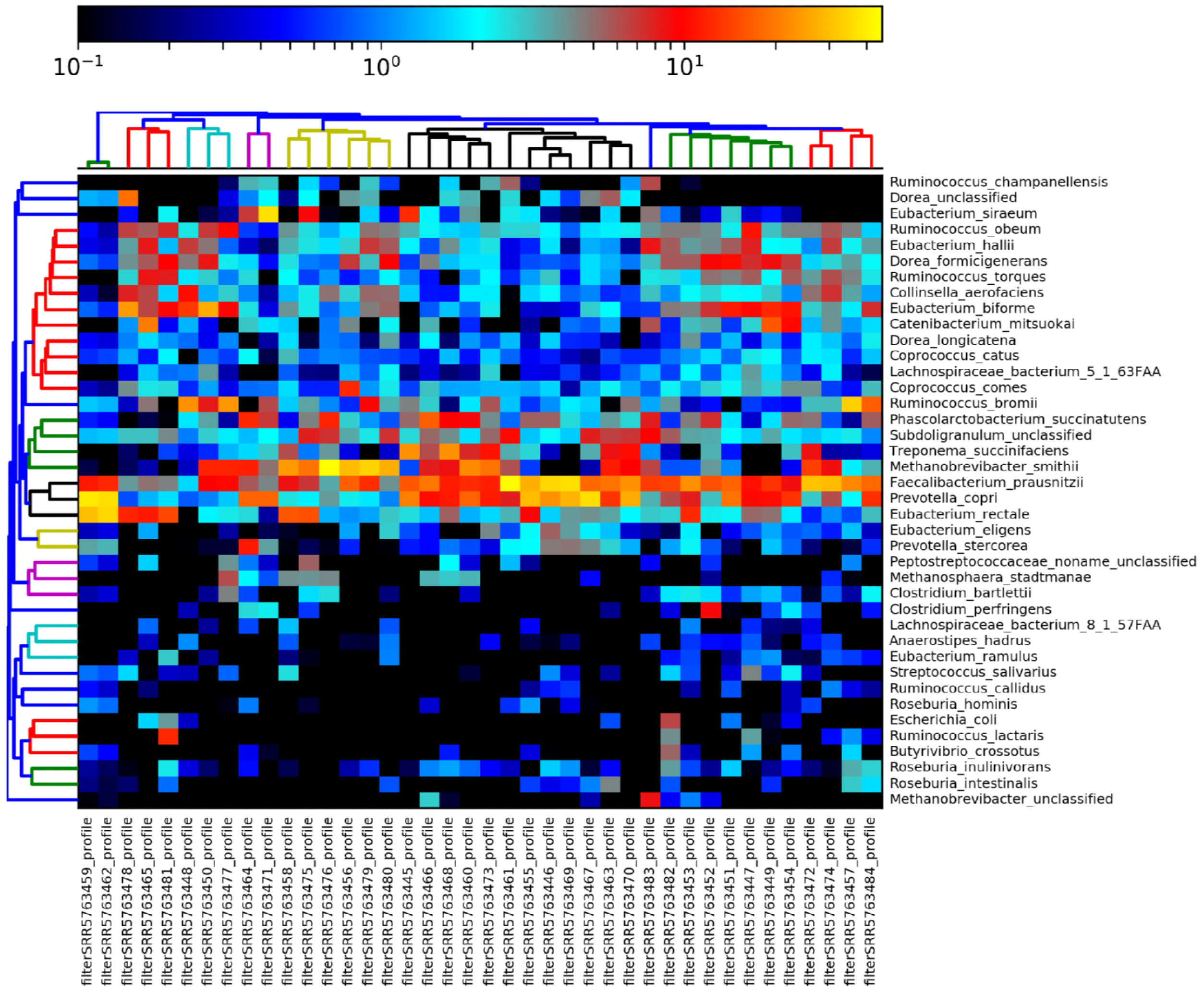
Heatmap showing Hadza gut microbial community composition. Drawn by MetaPhlAn2 based on species-abundance table where one row denotes a species and one column denotes a sample. Only 40 species with higher abundance are shown here. Main species in Hadza gut include *Faecalibacterium prausnitzii*, *Prevotella copri*, *Methanobrevibacter smithii*, *Eubacterium biforme*, which are represented in brighter-color cells.

From 116 species, we selected 33 species which existed in at least 8 samples in at least one season, to analyze their abundance differences across seasons. Of these 33 species, we identified 12 species whose abundance show significant (P<0.05, Wilcoxon) difference between dry and wet seasons, while insignificant difference between adjacent dry seasons. Their abundance distribution across seasons are shown in **Figure 2**. The Wilcoxon rank sum test results for all 33 species are shown in **Supplementary Table 1**.

**Figure 2.**
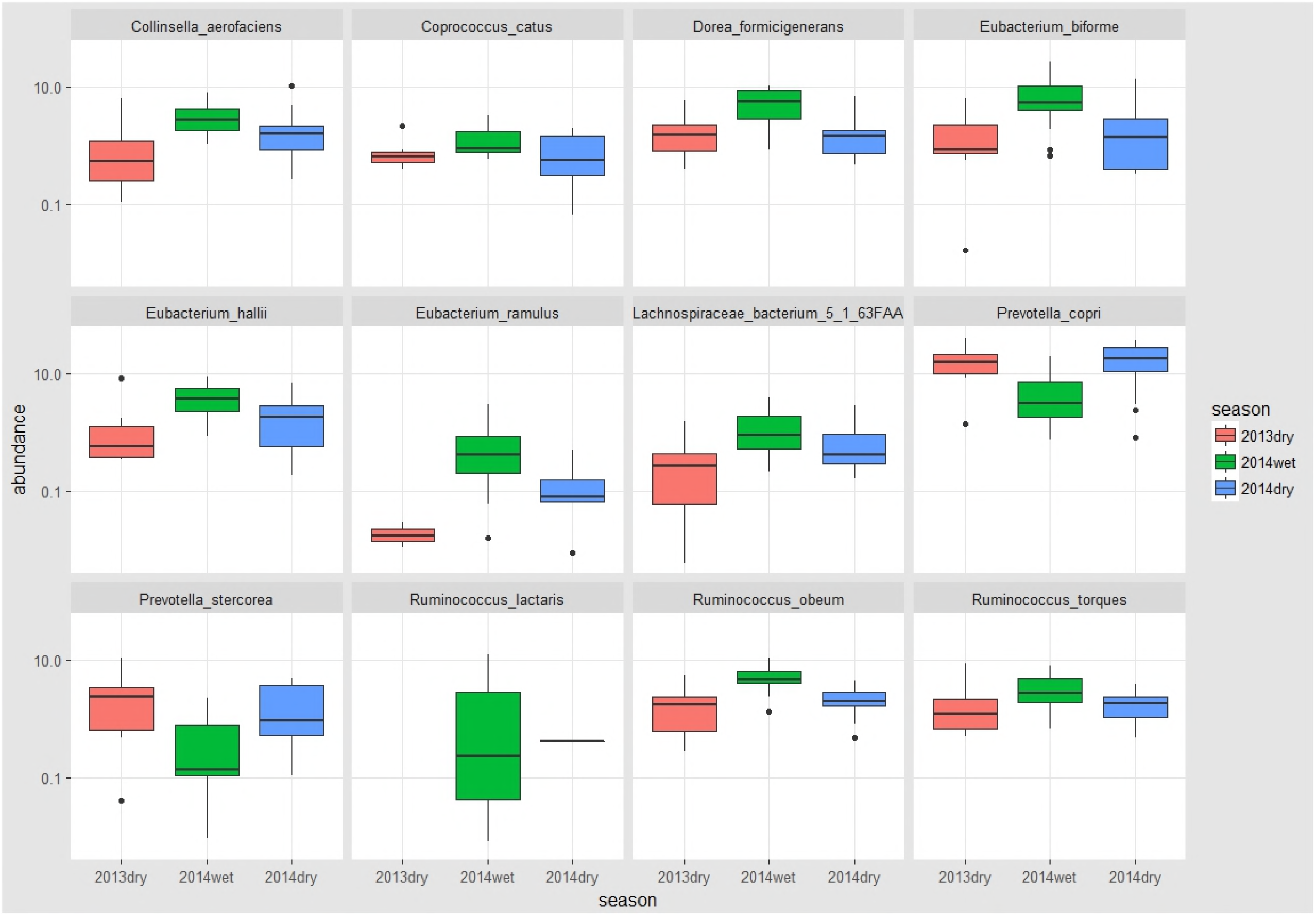
Abundance in certain gut microbial species are cyclic and differentiable by seasons. The changes in abundance across different seasons are shown with each panel representing one species. 2013dry samples (n =8), 2014wet samples (n = 19), 2014dry samples (n = 13) are denoted by red, green and blue, respectively. These 12 species have different (P<0.05, Wilcoxon) abundance between wet and dry seasons but indistinct (P>0.05, Wilcoxon) abundance between adjacent dry seasons.

Of these 12 species, only *Prevotella copri* and *Prevotella stercorea* have decreased abundance in 2014wet, which accords with the previous report indicating less *Prevotellaceae* in 2014wet[5]. *P.copri* has been proved to act in glucose metabolism and dietary high fiber can increase its abundance[19], which implies Hadza may have less fiber metabolism in wet seasons given the fact of all-year-around fiber consumption. All other 10 species show higher abundance in 2014wet, including *R.obeum* and *R.lactaris*, which all belong to Firmicutes. Since a previous report has found that Firmicutes show relatively stable abundance across seasons[5], the result here indicates that different species in Firmicutes may offset the difference.

### Strain-level SNP enrichment with the succession of seasons

Using the criteria that requires the presence of SNP in at least 3 samples, we select 55 prevalent species from 116 species to be included in the reference genome set. Performing the variant-calling procedure (see **Materials and Methods**), we used VarScan2 to identify SNP with a filtering threshold: minimum sequencing depth is 10 and minimum sequencing quality is 15. In general, we identified 765,106 SNP (8 samples, avg. 95,638) in 2013dry samples, 3,647,990 SNP (19 samples, avg 191,999) in 2014wet samples and 1,892,342 SNP (13 samples, avg. 145,564) in 2014dry samples. This demonstrates that there exists more SNP in wet season, and motivates us to investigate where these genome variations mainly originated.

15 species, whose average sequencing depth were above 10 in at least 3 samples, were selected for further analysis. Computing their SNP density (or namely SNP frequency,) in each sample, we found that all species show indistinct SNP density between dry seasons while 8 of them show enrichment (P<0.05, Wilcoxon) of SNP in wet season. These 8 species are *A. hadrus, C.mitsuokai, C.comes, E.biforme, E.hallii, R.inulinivorans, R.bromii* and *R.obeum*. SNP density distribution for all the species across seasons are shown in **Figure 3**. Wilcoxon rank sum test results for all the species are shown in **Supplementary Table 2**.

**Figure 3.**
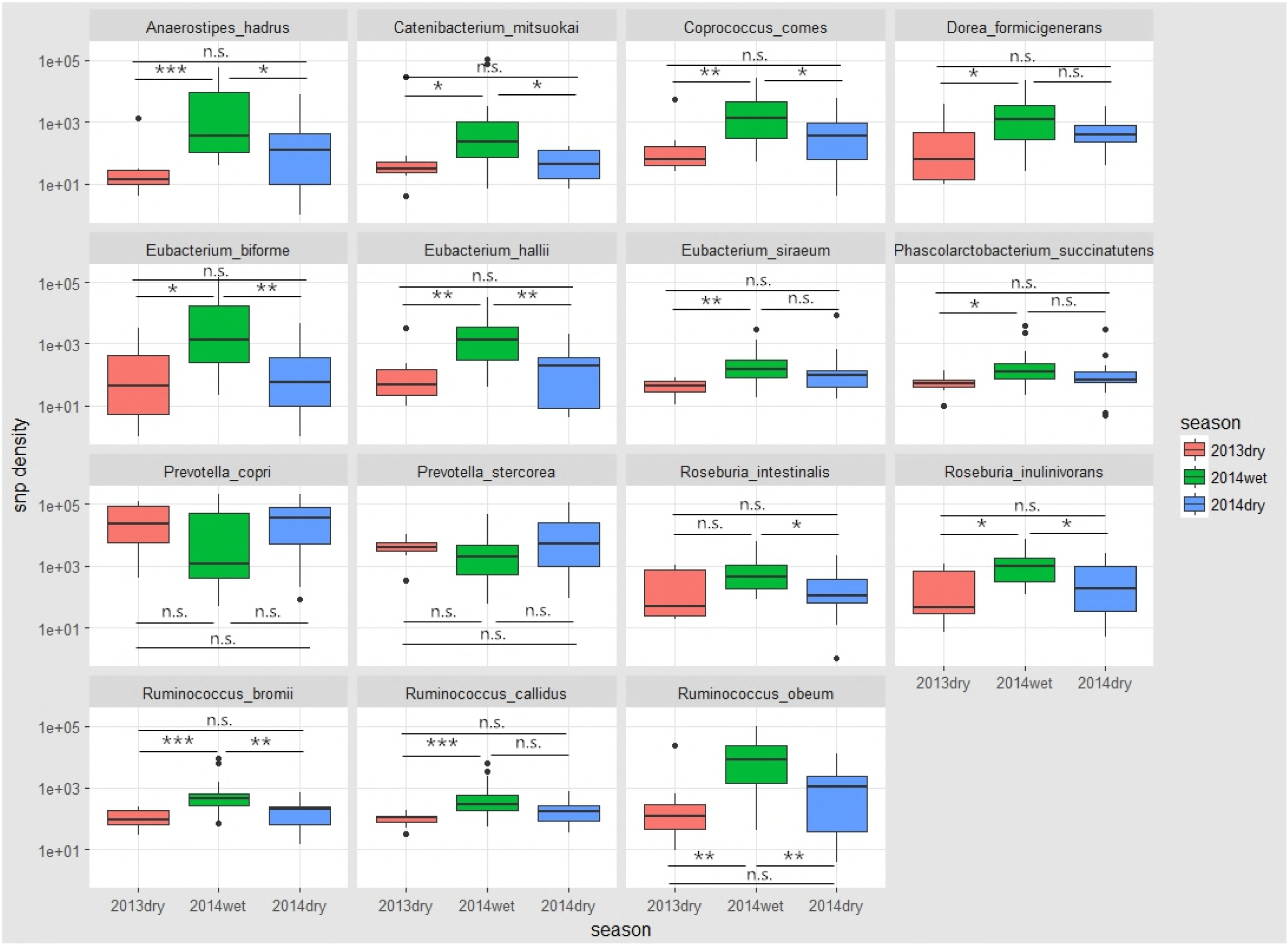
Certain species in Hadza gut microbiome have shown enrichment of SNPs in wet season. SNP density changes across seasons are shown with each panel representing one species. 8 of the 15 prevalent species have more SNP in wet season but comparable SNP in adjacent dry seaons. All boxplot distributions are tested by Wilcoxon rank sum test. Center values indicate the median; *P < 0.05, **P < 0.01, ***P<0.001; n.s., not significant.

The enrichment of SNPs in these species are highly correlated with the environmental and dietary changes accompanying the seasonal shifts. For example, it has been reported that dietary carbohydrate increases *A.hadrus*’s ability to produce butyrate[20] and dietary high protein decreases *R.bromii*’s metabolic ability[20]. Therefore, the enrichment of SNP in these species, which suggests more genomic variation, may correlate with the change in diet in different seasons.

Moreover, 5 out of the 8 species have indistinct abundance across seasons, which indicates that changes in species abundances may not be necessarily related to genomic variation. In other words, SNP characteristics are independent of species abundances, thus SNP characteristics would provide us with new ways for studying microbial community dynamics.

For each of the eight identified species, RAxML[21] was used to construct phylogenic trees based on whole genome SNP sites with mutated allele frequency bigger than 0.5 (see **Materials and Methods**). The best tree was visualized by R package ggtree. Two examples for species *E.biforme* and *E.hallii* are shown in **Figure 4**. Most wet-season samples are in one cluster and dry-season samples are in the other cluster, which indicates the generation of new strain in wet season samples. The trees for remaining 6 species are in **Supplementary Figure 1**. Interestingly, we found that one sample (SRA run: SRR5763465) is special: This dry-season sample is mixed with other wet-season samples. From its subject information, we found this sample possibly came from a kid less than 3 years old. The environmental stressors and dietary patterns of this host might be different from other members of the Hadza people.

**Figure 4.**
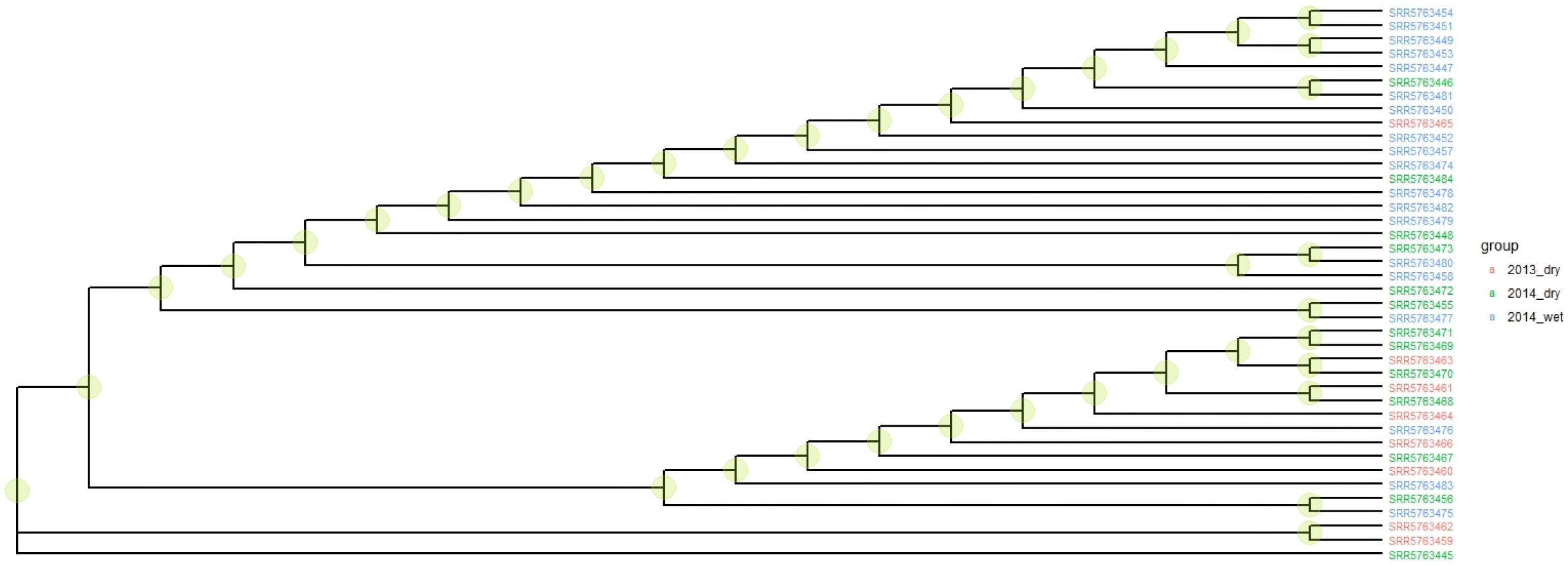
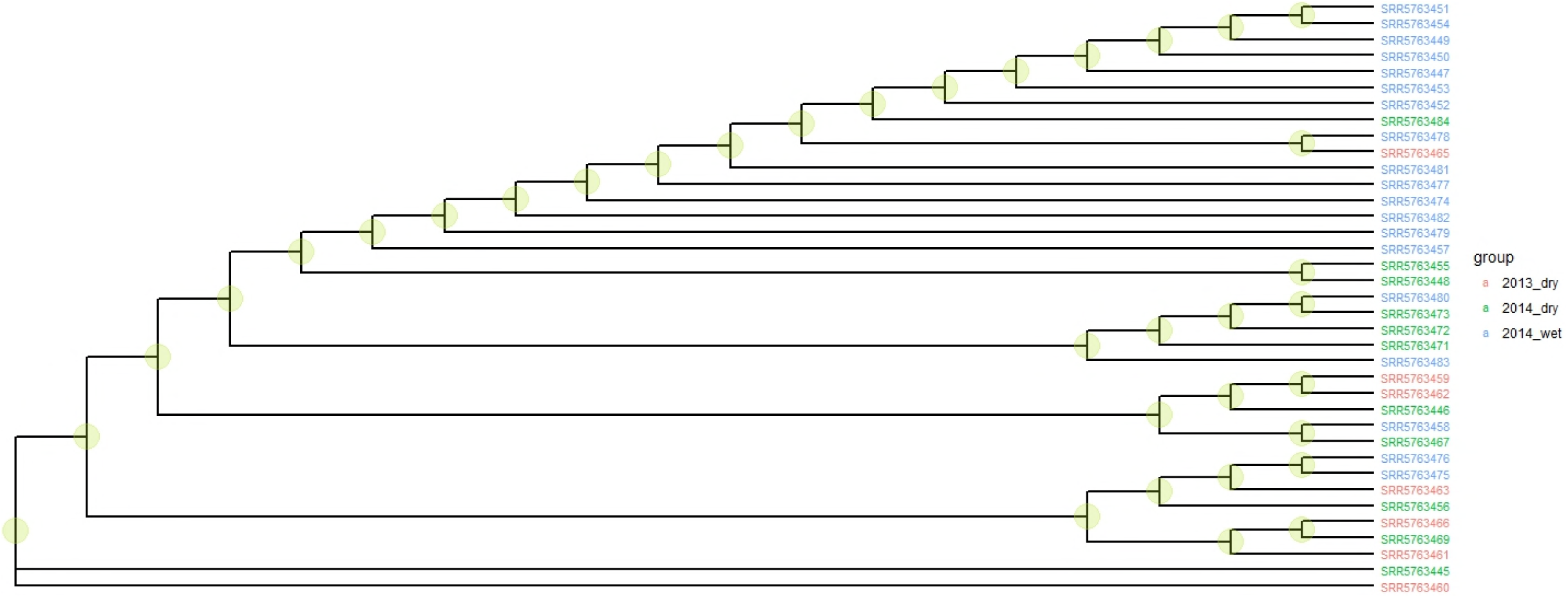
Wet-season samples are separable from dry-season samples in phylogenic trees. Phylogenic trees based on whole genome SNP sites for two species are shown here. Each branch represents one sample and samples from different seasons are differentiated by three colors. Clear clustering is observed in both trees where wet-season samples are restricted in one cluster and dry-season samples in the other. In **(a)** *E.hallii*(top panel), the top cluster contains 16 wet-season samples(84.21% of total wet-season samples). The bottom cluster contains 14 dry-season samples(66.67% of total dry season samples). Similarly in **(b)** *E.biform*(bottom panel), the top cluster contains 14 wet-season samples(73.68%). The bottom cluster contains 14 dry-season samples(77.78% of total dry season samples).

In some specific areas of the phylogenic trees, we can observe the mingling of wet-season samples and dry-season samples, which may be attributed to the inclusion of samples from kids and repeated sampling in single season. Actually, these 5 samples were excluded in Samuel A. Smits’s analysis. Also, the limited number of samples may enlarge the bias in single sample and add difficulty to the discovery of general conclusions.

We also extracted SNP sites whose mutated allele frequency are larger than 0.2 in each sample to obtain a matrix used for calculating pairwise distance between samples. The dist function in R was used with the parameter: method = ‘manhattan’. The hclust function was then used with the parameter: method = ‘complete’ to perform hierarchical clustering on all the samples. The results for species *E.biforme* and *E.hallii*, visualized in ggtree, are shown in **Figure 5** as examples. The results for the remaining six species are in **Supplementary Figure 2**. Similar to the previous phylogenic tree result, we found wet season samples are separable from dry season samples.

**Figure 5.**
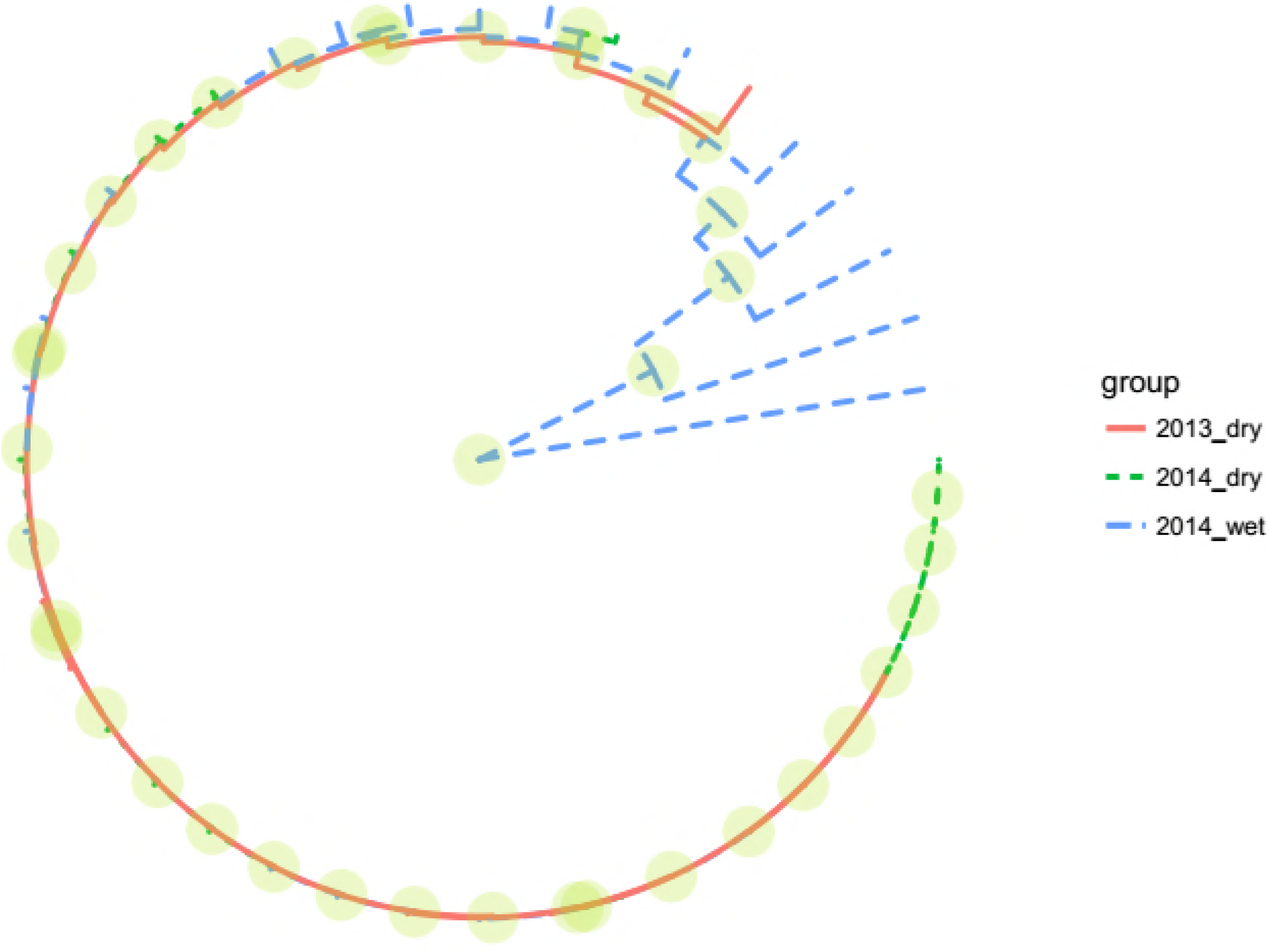
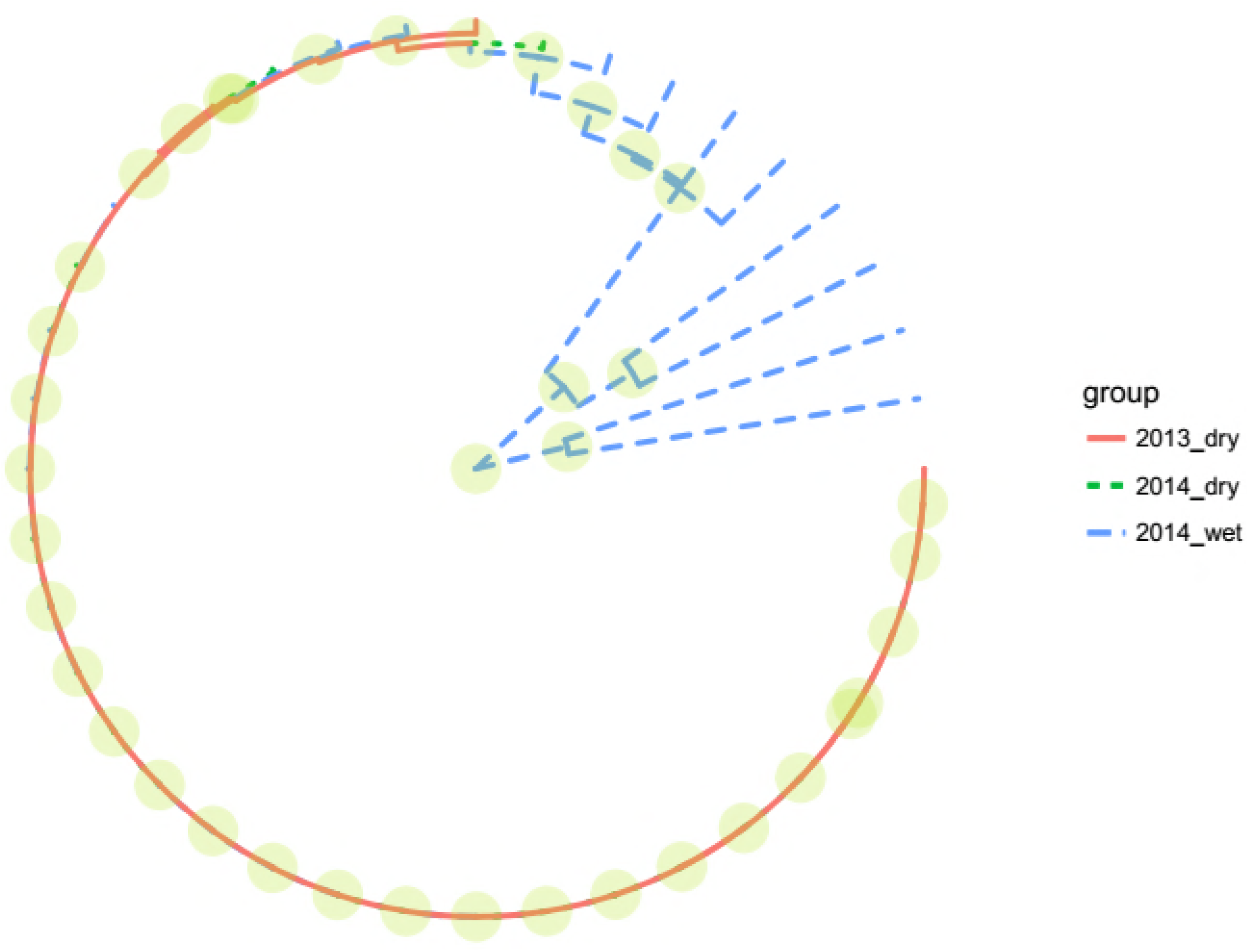
Wet-season samples are separable from dry-season samples based on their SNP profiles. Clustering results based on whole genome SNP sites for two species are shown here. Each branch represents one sample and samples from different seasons are differentiated by three colors. Clear clustering is observed in **(a)** *E.hallii*(left panel) and **(b)** *E.biform*(right panel) where wet-season samples are restricted in one cluster and dry-season samples in the other, which indicates the generation of new strain in 2014wet.

### Gene-level SNP enrichment with the succession of seasons

From a total of 23,504 genes in the 15 selected species, 5,959 genes had SNP in at least 8 samples and were targeted for further analysis. We identified 827 genes whose SNP density showed significant difference (P<0.05, Wilcoxon) between wet season and dry season, but similar pattern between dry seasons. Among these 827 genes, 83 genes showed most significant difference (P<0.01, Wilcoxon). Information on the 83 genes is shown in **Supplementary Table 3**.

For each of the 83 genes, the protein sequence was used to search by BLAST against the KEGG annotation and the best hit was selected for its pathway information. 36 genes were annotated to 52 KEGG pathways (**Supplementary Table 4**) and pathways that involve at least two genes were selected to show in **Figure 6**. Many of these genes are from *Ruminococcus obeum*, concentrating on metabolic pathways like carbon metabolism, pyruvate metabolism and glycolysis, as shown by KEGG annotation. This implies that the seasonal changes might indirectly impact the mutational patterns for specific species and functions for gut microbiome of an unindustrialized population, indicating the role of these variants in their adaptation to the changing environment and diets.

**Figure 6.**
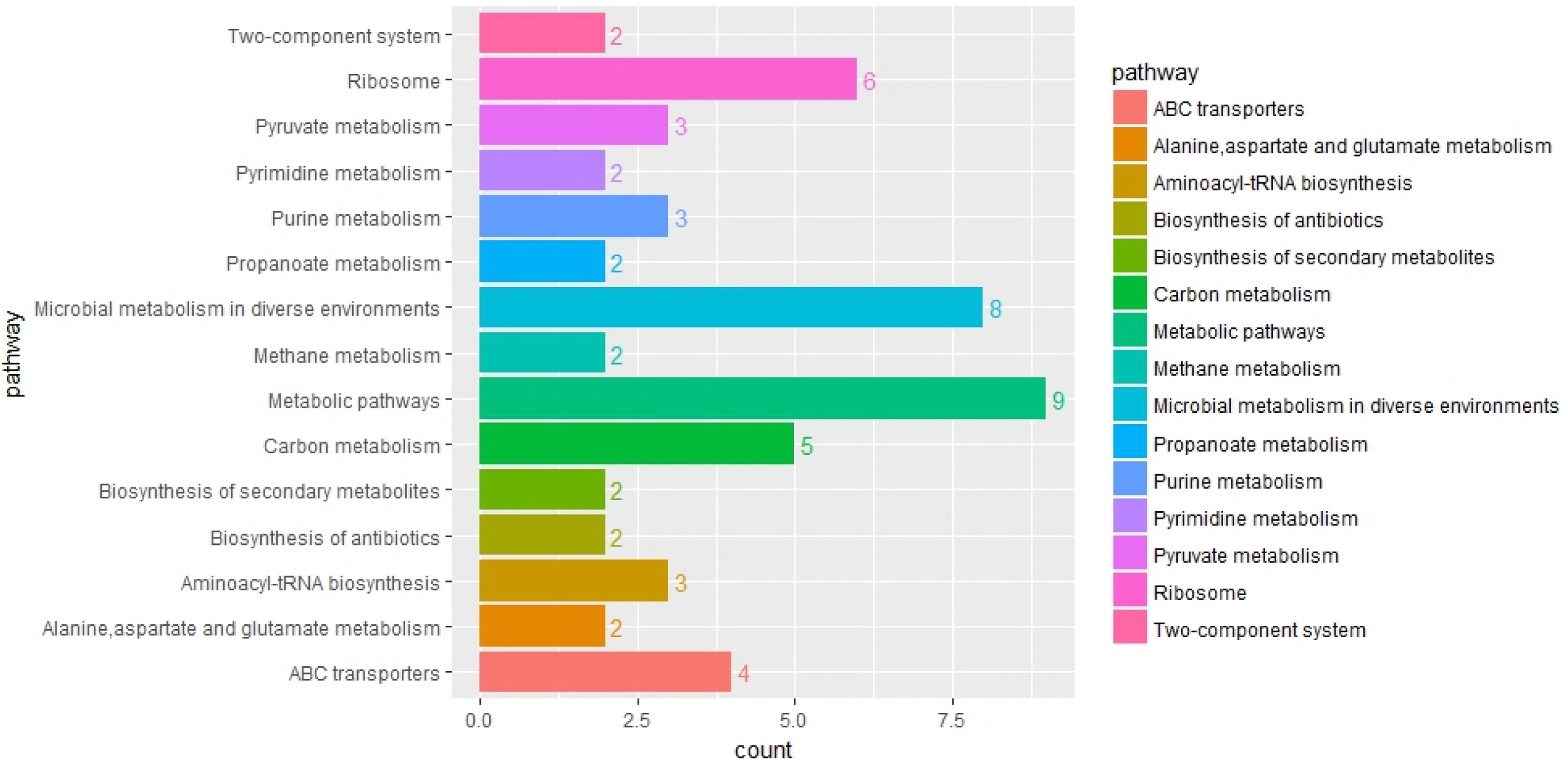
Pathway histogram of genes with different SNP pattern across seasons. Each bar represents a specific biological pathway and the number above it denotes the number of genes involved in this pathway. In Hadza gut microbiome, the genes with different SNP density(P<0.01, Wilcoxon) in 2014wet mainly participate in metabolic pathways like carbon metabolism and ribosome.

## Conclusion

This work has implications from both technical and microbiological viewpoints. From a technical viewpoint, we identified that VarScan2 showed comparatively better performance for SNP calling from metagenomic dataset, in both selectivity and sensitivity. Thus, we recommend using VarScan2 with filtering in human gut metagenome analysis. This analysis provides a reference for future investigation in variant calling and suggests that the direction to improve variant-calling tools is to improve the identification efficiency and reduce false positive results at the same time.

From microbiological viewpoint, we have identified the characteristic enrichment of SNPs in species and functions with the succession of seasons. Strain-level SNP analysis of Hadza gut metagenome identified eight species with significant enrichment of SNP in wet season. Phylogenic trees and hierarchical clustering results of their whole genome SNP both demonstrate the emergence of new strains in wet season. Eight prevalent species have significant SNP enrichments in wet season of which only three species have relatively high abundances. This indicates that SNP characteristics are independent of species abundances, and would provide new ways to study microbial community dynamics. Additionally, we identify 83 genes with most significantly characteristic SNP distributions between wet season and dry season. Many of these genes are from *Ruminococcus obeum*, concentrating on metabolic pathways like carbon metabolism, pyruvate metabolism and glycolysis, as shown by KEGG annotation. This implies that the seasonal changes might indirectly impact the mutational patterns for specific species and functions for gut microbiome of an unindustrialized population, indicating the role of these variants in their adaptation to the changing environment and diets.

The results here unveil the species-level and genome-level variations in gut microbiome along the succession of seasons, clarifying the role of these variants in their adaptation to the changing environment and diets. These have also laid a foundation for future investigation of gut microbiome dynamics in response to changes in lifestyle and other environmental stressors.

## Materials and Methods

### Simulated and real datasets

Simulated data were generated by three steps: We first generated a SNP list containing 8,148 sites from 4 species residing in human gut(*Bacteroides caccae*, *Bacteroides fragilis*, *Bacteroides ovatus*, *Bacteroides vulgatus*). Then, a mutated reference genome set was achieved by replacing corresponding SNP sites. BBMap was applied to generate 1,000,000 paired-end simulated reads based on the mutated reference genome set with the parameters: SNPrate=0.002 insrate=0.00002 delrate=0.00002, which specified the simulated sequencing error.

Hadza gut microbiome dataset were whole genome sequencing data that are described in [5], all of the 40 metagenomic samples spanned four subseasons (2013 Dry, 2014 Early Wet, 2014 Late Wet, 2014 Dry). All of these microbiome data were downloaded from NCBI SRA database (accession SRA582120).

### SNP calling by bcftools, GATK and VarScan2

The processed bam files were used for SNP calling by three tools. The command for BCFtools was “samtools mpileup -ugf genome_for_sim.fna dupfree_sim_meta.bam | bcftools call -vmO v -V indels -o bcftools_meta.vcf”, where smtools -u specified uncompressed file, -g specified output as bcf format, -f specified reference sequence file, BCFtools -v specified only output variant sites, -O v specified output file as uncompressed vcf format and -V indels specified ignoring indel.

For GATK, SAMtools faidx and picard CreateSequenceDictionary were used to generate .fai file and .dict file, respectively to index the reference file. The command is “java -jar GenomeAnalysisTK.jar -T HaplotypeCaller -R genome_for_sim.fna -I dupfree_sim_meta.bam -o GATK_meta.vcf”.

For VarScan2, samtools mpileup was first used to output mpileup file and the command is:

java -jar VarScan.v2.3.9.jar mpileup2SNP sim_meta.mpileup --min-coverage 1 --output-vcf 1--variants --min-reads2 1 --min-avg-qual 1 > varscan_meta.vcf, where --min-coverage specified the minimum sequencing depth,--variants specified only output variants,--min-reads2 specified the minimum number of reads that support the SNP, --min-avg-qual specified the minimum sequencing quality.

### Sensitivity and selectivity calculating

Comparing SNP identified by the three tools with actual SNP (ground truth), we assessed their performance from two aspects which were calculated as follows.

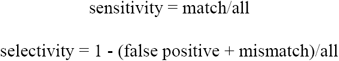

Here, ‘match’ represents the number of actual SNP that are identified by the tool. ‘false positive’ represents the number of SNP identified that are actually non-existent. ’mismatch’ represents the number of actual SNP that are identified as wrong base by the tool. ‘all’ represents the number of all actual SNP.

### Variant calling procedure

Clean reads of 40 samples were aligned to reference genome set via BWA MEM with the parameter -R to specify the header group in output sam files. SAMtools view command was used with parameter -bS to transform bam files into bam files. SAMtools sort command was used to sort bam files by the order of chromosome, which is actually scaffold here. Picard MarkDuplicates command was used with parameter REMOVE_DUPLICATES=true to remove pcr duplicates. Samtools mpileup command was used with parameter -Bf to specify reference genome set and output mpileup files. VarScan mpileup2SNP command was used with parameter --min-coverage 10 --output-vcf 1 --variants --min-avg-qual 15 to identify SNP from mpileup files and output vcf files. The variant calling procedure was illustrated in **Figure 7**.

**Figure 7.**
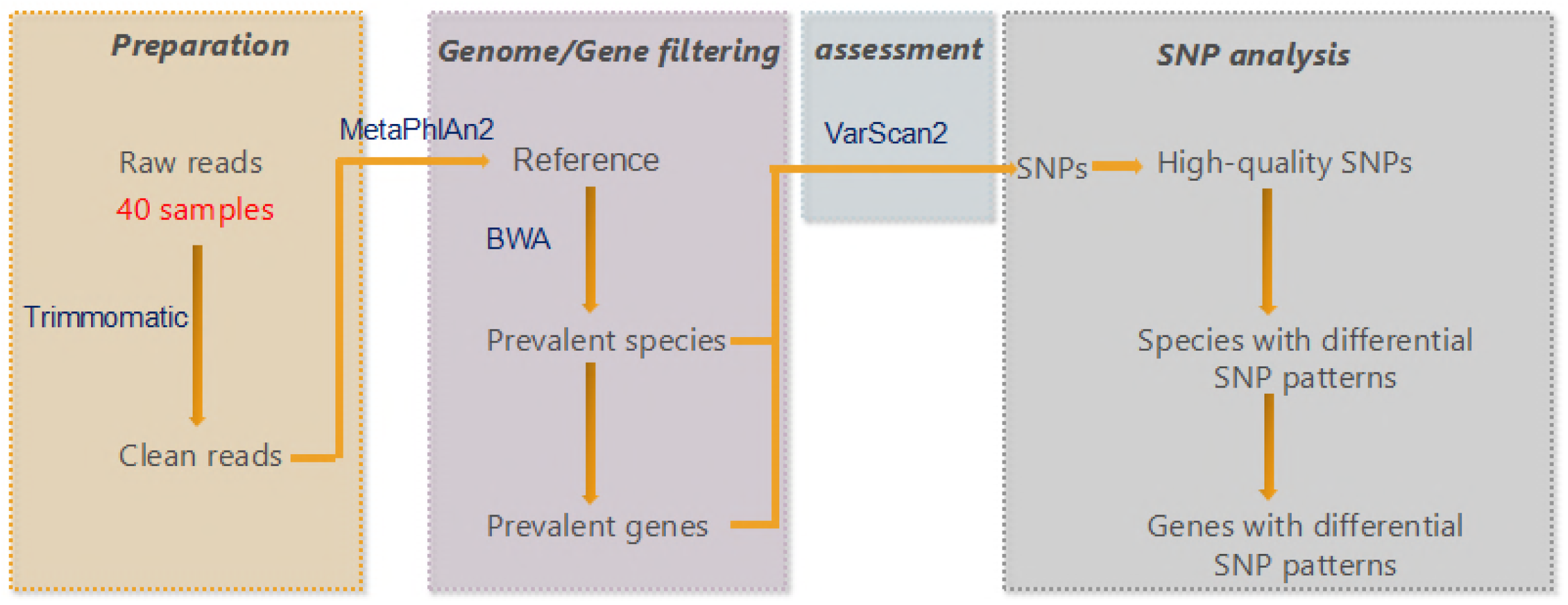
Variant calling procedure. Each colored block represented a series of analysis for data preparation, genome/gene filtration and SNP analysis.

### Phylogenic tree construction

The parameters used for RAxML to construct phylogenic trees based on whole-genome SNP sites were -m ASC_GTRGAMMA -p 12345 --asc-corr=lewis -f a -x 12345 -#100, which specified the nucleotide substitution model using -m, the ascertainment bias correction method using –asc-corr, and bootstrap numbers using -#.

## Acknowledgments

We thank Kal Ng from University of Malaya for discussion about the topic, as well as proof-reading. This work is partially supported by National Science Foundation of China grant 31871334 and 31671374, Ministry of Science and Technology’s high-tech (863) grant 2014AA021502, and Sino-German Research Center grant GZ878.

## Competing financial interests

The authors declare no competing financial interests.

## References

1. Clemente, J.C., et al., The impact of the gut microbiota on human health: an integrative view. Cell, 2012. 148(6): p. 1258–70.

2. Kundu, P., et al., Our Gut Microbiome: The Evolving Inner Self. Cell, 2017. 171(7): p. 1481–1493.

3. Ehlers, S., S.H. Kaufmann, and C. Participants of the 99 Dahlem, Infection, inflammation, and chronic diseases: consequences of a modern lifestyle. Trends Immunol, 2010. 31(5): p. 184–90.

4. Schnorr, S.L., et al., Gut microbiome of the Hadza hunter-gatherers. Nat Commun, 2014. 5: p. 3654.

5. Smits, S.A., et al., Seasonal cycling in the gut microbiome of the Hadza hunter-gatherers of Tanzania. Science, 2017. 357(6353): p. 802–806.

6. Bagel, S., et al., Impact of gyrA and parC mutations on quinolone resistance, doubling time, and supercoiling degree of Escherichia coli. Antimicrob Agents Chemother, 1999. 43(4): p. 868–75.

7. Sokurenko, E.V., et al., Pathogenic adaptation of Escherichia coli by natural variation of the FimH adhesin. Proc Natl Acad Sci U S A, 1998. 95(15): p. 8922–6.

8. Rocap, G., et al., Genome divergence in two Prochlorococcus ecotypes reflects oceanic niche differentiation. Nature, 2003. 424(6952): p. 1042–7.

9. Schloissnig, S., et al., Genomic variation landscape of the human gut microbiome. Nature, 2013. 493(7430): p. 45–50.

10. Zhu, A., et al., Inter-individual differences in the gene content of human gut bacterial species. Genome Biol, 2015. 16: p. 82.

11. Chen, Y., et al., Gut metagenomes of type 2 diabetic patients have characteristic single-nucleotide polymorphism distribution in Bacteroides coprocola. Microbiome, 2017. 5(1): p. 15.

12. McKenna, A., et al., The Genome Analysis Toolkit: a MapReduce framework for analyzing next-generation DNA sequencing data. Genome Res, 2010. 20(9): p. 1297–303.

13. Storey, J.D. and R. Tibshirani, Statistical significance for genomewide studies. Proc Natl Acad Sci U S A, 2003. 100(16): p. 9440–5.

14. Koboldt, D.C., et al., VarScan 2: somatic mutation and copy number alteration discovery in cancer by exome sequencing. Genome Res, 2012. 22(3): p. 568–76.

15. Li, H., et al., The Sequence Alignment/Map format and SAMtools. Bioinformatics, 2009. 25(16): p. 2078–9.

16. Bolger, A.M., M. Lohse, and B. Usadel, Trimmomatic: a flexible trimmer for Illumina sequence data. Bioinformatics, 2014. 30(15): p. 2114–20.

17. Truong, D.T., et al., MetaPhlAn2 for enhanced metagenomic taxonomic profiling. Nat Methods, 2015. 12(10): p. 902–3.

18. Han, C., et al., Complete genome sequence of Treponema succinifaciens type strain (6091). Stand Genomic Sci, 2011. 4(3): p. 361–70.

19. Kovatcheva-Datchary, P., et al., Dietary Fiber-Induced Improvement in Glucose Metabolism Is Associated with Increased Abundance of Prevotella. Cell Metab, 2015. 22(6): p. 971–82.

20. Mu, C., et al., Temporal microbiota changes of high-protein diet intake in a rat model. Anaerobe, 2017. 47: p. 218–225.

21. Stamatakis, A., RAxML version 8: a tool for phylogenetic analysis and post–analysis of large phylogenies. Bioinformatics, 2014. 30 (9): p. 1312–3.

